# Evaluation of the impact of seasonal variations in photoperiod on the hepatic metabolism of medaka (*Oryzias latipes*)

**DOI:** 10.1101/745646

**Authors:** Koichi Fujisawa, Taro Takami, Haruko Shintani, Nanami Sasai, Toshihiko Matsumoto, Naoki Yamamoto, Isao Sakaida

## Abstract

Organisms living in temperate regions are sensitive to seasonal variations in the environment; they are known to accumulate energy as fat in their livers during the winter when days are shorter, temperatures are lower, and food is scarce. However, the impact of variations in photoperiod alone on hepatic lipid metabolism has not been well-studied. Therefore, in this study, we analyzed lipid metabolism in the liver of medaka, *Oryzias latipes*, while varying the length of days at constant temperature. Larger amounts of fatty acids accumulated in the liver after 14 days under short-day conditions than under long-day conditions. Metabolome analysis showed no accumulation of the long-chain unsaturated fatty acids required at low temperatures, but showed a significant accumulation of long-chain saturated fatty acids. Short-day conditions induced decreased levels of succinate, fumarate, and malate in the tricarboxylic acid cycle, decreased expression of PPARα, and decreased accumulation of acylcarnitine, which suggested inhibition of lipolysis. In addition, when a high-fat diet was administered to transparent medaka under short-day conditions, larger amounts of fat accumulated and medaka with fatty liver were efficiently produced. Detailed analysis of the relationship between seasonal changes and hepatic steatosis will be important in the future as hepatic diseases become more prevalent in modern society; the findings obtained in our study will be useful for research studies pertaining to the relationship between photoperiod and disorders such as hepatic steatosis and non-alcoholic steatohepatitis.

## Introduction

Lifestyle habits such as diet, exercise, rest, as well as smoking and drinking, play a role in the onset and progression of lifestyle-related disease such as diabetes, obesity, hyperlipidemia and hypertension. Among lifestyle-related diseases, fatty liver caused by overnutrition is the most frequently encountered liver disease, and the number of patients with the condition continues to increase. Fatty liver disease is a general term for disorders in which triglycerides are deposited in hepatocytes and cause liver damage; in individuals with no apparent history of alcohol consumption, the condition is known as nonalcoholic fatty liver disease (NAFLD). NAFLD can be classified into two different types: simple fatty liver which has a benign prognosis, and nonalcoholic steatohepatitis (NASH), which is a progressive condition. NASH develops into cirrhosis and liver cancer (Lapadat et al., 2017). Thus far, mice have been used to develop methods aimed at inhibiting the progression of fatty liver disease, but new and more efficient models have been desired (Asgharpour et al., 2016).

Compared to rodents such as mice, small fish species such as medaka (*Oryzias latipes*) and zebrafish (*Danio rerio*) have recently attracted attention as new model organisms because of their smaller size, which can help reduce rearing space and costs. Their advantages also include their completed genome and established methods for the preparation of transgenic and knockout fish; they are highly fertile and take less time to reach maturity; and large-scale screenings are easy to perform. Further, among small fish species, medaka are capable of living through the winter, are said to have the same glucose and lipid metabolic functions found in mammals, and are highly capable of accumulating fat in their livers.

We previously reported the utility of a medaka model with a high-fat diet (HFD) for tissue analysis and lipid analysis (Matsumoto et al., 2010). In addition, we evaluated HFD-induced hepatic steatosis through direct observation and ultrasound of transparent medaka for non-invasive analyses of the progression of HFD-induced fatty liver disease (Fujisawa et al., 2018). Further, direct observation and ultrasound examination of transparent medaka allowed for the evaluation of hepatic steatosis induced by the administration of alcohol (Fujisawa et al., 2019).

Organisms which carry out life activities under the influence of seasonal cycles are known to display changes in various physiological functions related to photoperiodism, such as their breeding activities. In birds and mammals, the pars tuberalis (PT) of the adenohypophysis, which is located at the base of the pituitary gland, is known to play a central role by regulating the hypothalamo-hypophyseal system through secretion of various signal and functional molecules (Ikegami and Yoshimura, 2012). In addition, fat accumulation in the inguinal region, epididymis and retroperitoneal region of Japanese grass voles has been previously related to the effect of light on living organisms (Dark and Zucker, 1986). In fish, recent reports on light-sensing systems have shown that coronet cells present in the saccus vasculosus of the masu salmon (*Oncorhynchus masou*) act as “seasonal sensors” by sensing changes in the number of daylight hours, and control breeding activities (Nakane et al., 2013).

Thus far, there have been no detailed reports on the length of photoperiod and liver metabolism in small fish species. Therefore, in this study, we assessed the types of changes occurring in the liver of medaka under long- and short-day conditions. In addition, we examined whether an experimental fatty liver could be induced more efficiently by varying the length of daylight in a breeding room with a conventional photoperiod of light:dark ratios of 14:10 or 12:12 for ovum collection.

## Results

### Fat accumulates in large amounts in the liver under short day conditions

After the 14/10 long-day conditions (L:D=14:10), evaluations were conducted for a 20-day period under 10/14 short-day conditions (L:D=10:14). Findings regarding the medakas’ amount of activity confirmed that immediately after breeding conditions shifted to short-day conditions, the duration of activity changed to 10 h (Fig.1A).

**Fig. 1.**
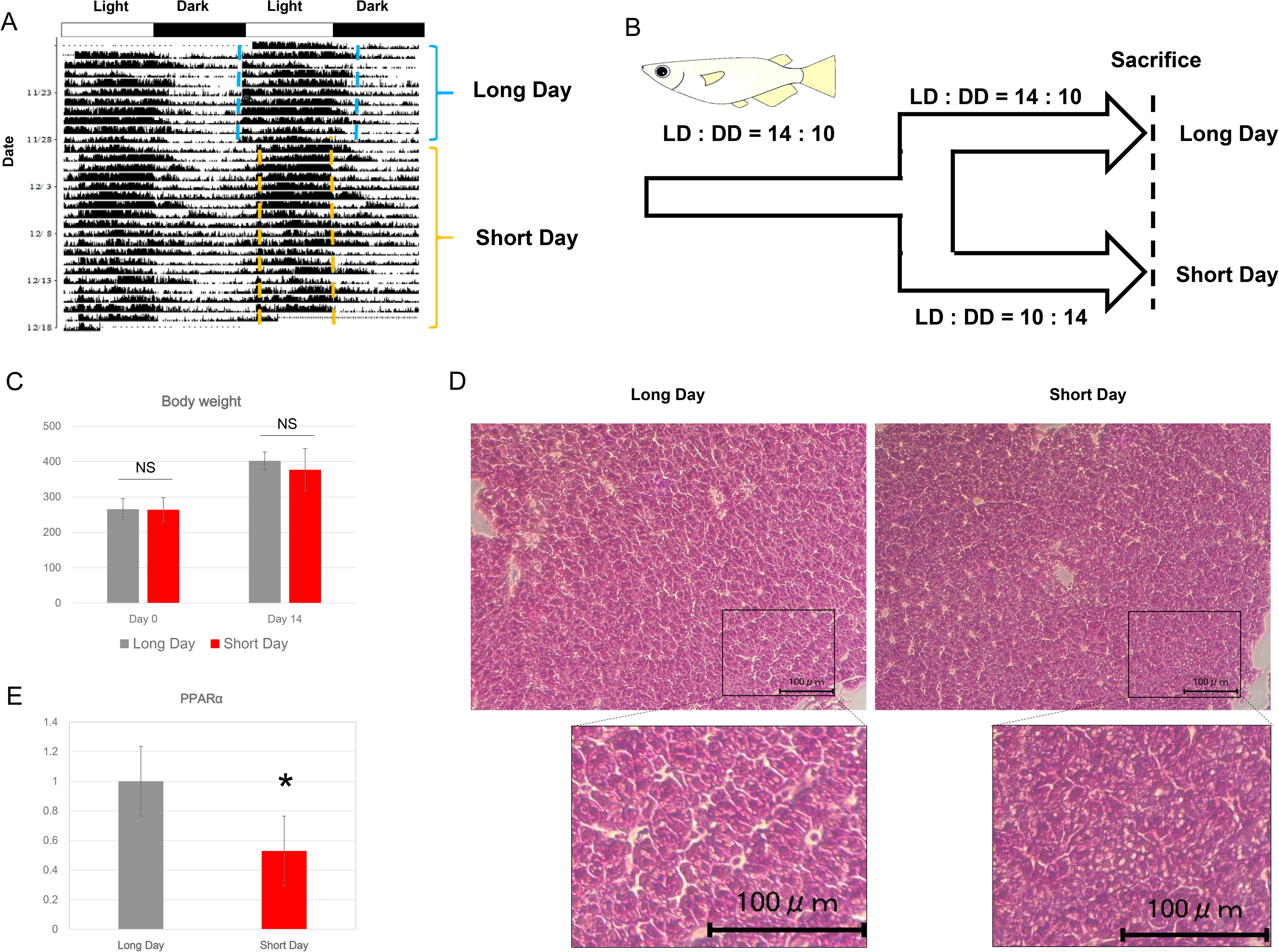
Photoperiod-dependent changes in the liver of medaka fed with a normal diet. (A) Double-plotted actograms of typical medaka. The blue dotted line indicates long-day conditions (L:D=14:10), and the yellow dotted line indicates short-day conditions (L:D=10:14). (B) Schematic illustration of experiments conducted on medaka reared under long-day and short-day conditions. (C) Changes in the the medakas’ body weight at day 14 after exposure to variations in photoperiod. (D) Hematoxylin-eosin staining of medaka liver reared under long-day and short-day conditions. (E) Real-time PCR analysis of the expression of PPARα. EF1α was used as internal control. Data are represented as mean ± SD. * indicates significant difference (P ≤ 0.05)

Next, the effect of photoperiod on hepatic lipid metabolism was evaluated. The medakas’ body weight did not differ significantly depending on photoperiod (Fig.1B and C). Hematoxylin-Eosin (HE) staining of the liver showed that the amounts of lipid droplets were larger in the short-day group (Fig. 1D). Real-time PCR analysis showed that the expression of PPARα, a gene involved in lipid metabolism, had significantly decreased in the short-day group (Fig.1E).

### Long-chain fatty acids and acylcarnitine accumulate in the liver under short-day conditions

HE staining showed increased fat deposition in the short-day group; therefore, a more detailed analysis of metabolites was carried out through metabolomic and principal component analysis (PCA) of both the short-day and long-day groups (Fig.2A). Interestingly, levels of metabolites involved in the tricarboxylic acid cycle (TCA) cycle, such as succinate, malate, and fumarate (Fig.2B), showed a significant decrease. Meanwhile, analysis of metabolites involved in the glycolytic pathway showed no change except in the levels of glucose (Fig.2B). As for lipid-related changes under short-day conditions, increased levels of long-chain fatty acids such as palmitate (16:0), margarate (17:0), and stearate (18:0) (Table.1) were observed. Meanwhile, there was little change in polyunsaturated fatty acids (PUFA) levels, which had been reported to increase at low temperatures (Table 2). In addition, the amounts of several long-chain acylcarnitines had significantly increased (Table 3). IPA analysis showed that the glutathione (GSH) and taurine biosynthesis pathways were canonical pathways, and levels of GSH decreased while levels of taurine increased (Fig.3 A and B).

**Fig. 2.**
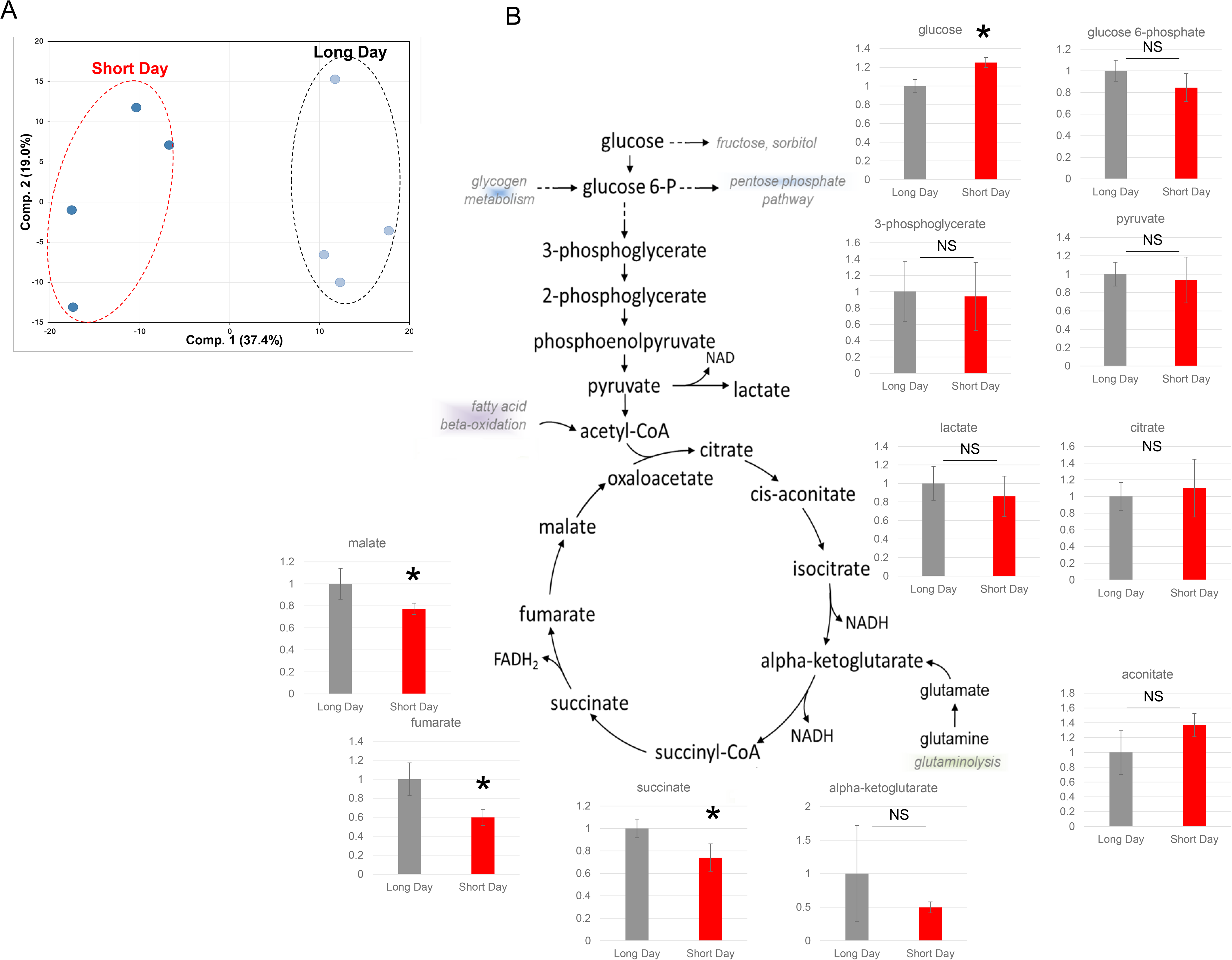
Metabolomic analysis of medaka liver. (A) Principal component analysis of normalized metabolic data obtained from the liver of medaka reared under long-day and short-day conditions. Percentage values indicated on the axes represent the contribution rate of the first (PC1) and second (PC2) principal components to the total amount of variation. Identical photoperiods are encircled by dotted lines of the same color. (B) Changes in metabolites involved in the glycolytic pathway and TCA cycle. The vertical axis represents relative peak area.

**Fig. 3.**
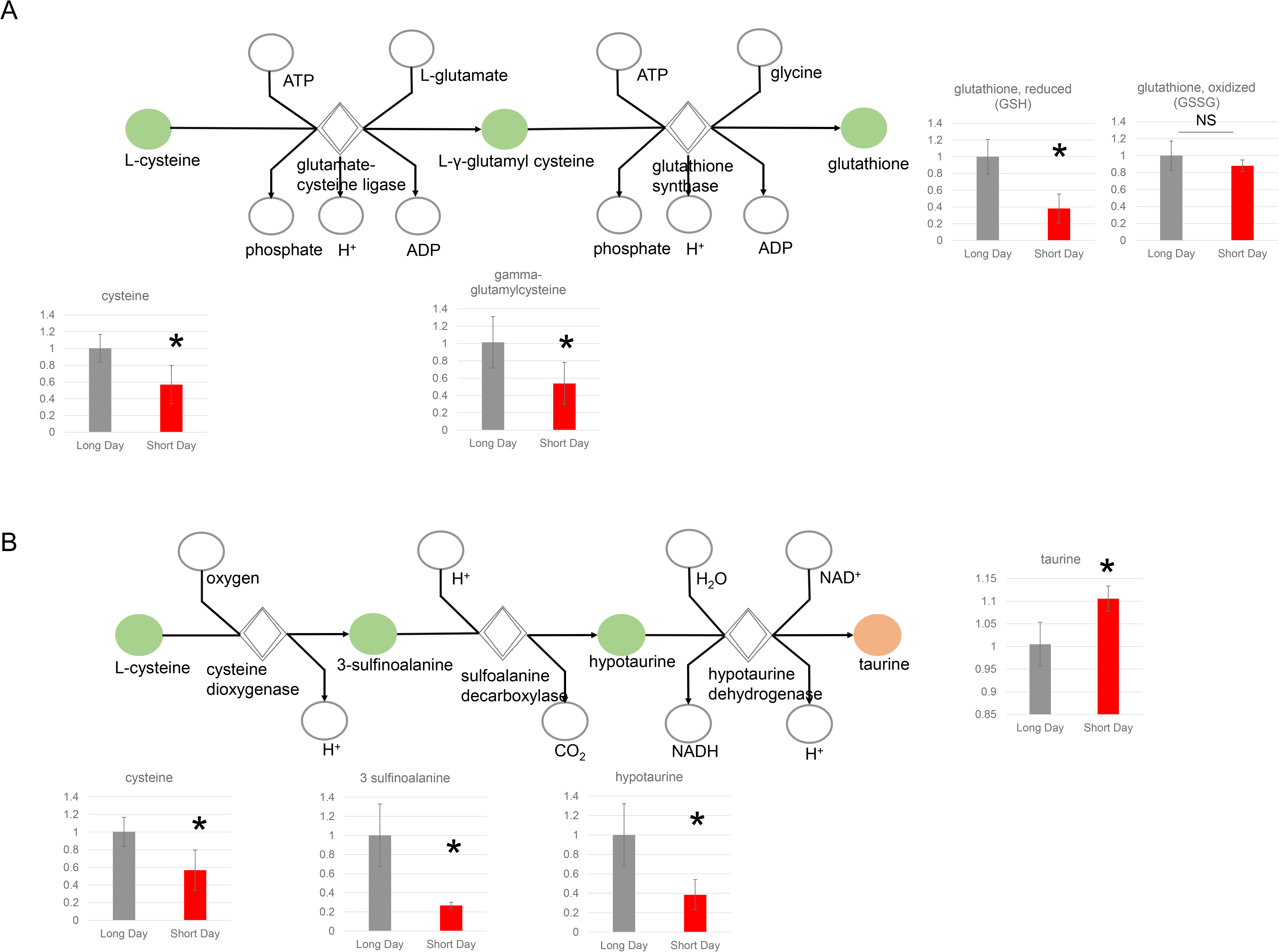
Metabolomic changes involved in glutathione biosynthesis and taurine biosynthesis. (A) Changes in metabolites involved in glutathione biosynthesis. (B) Changes in metabolites involved in taurine biosynthesis. Data are represented as mean ± SD. * indicates significant difference (P ≤ 0.05). Green color indicates downregulated metabolites. Orange color indicates upregulated metabolites.

**Table 1.** Changes in metabolites involved in long chain fatty acid metabolism. Red: indicates significant difference (P ≤ 0.05) between the groups shown; metabolite ratio of ≥ 1.00.

| Sub Pathway | Biochemical Name | Short day /Long day |
| --- | --- | --- |
| **Long Chain Fatty Acid** | myristate (14:0) | 1.51 |
|  | myristoleate (14:1n5) | 1.35 |
|  | pentadecanoate (15:0) | 2.10 |
|  | palmitate (16:0) | 1.57 |
|  | palmitoleate (16:1n7) | 1.61 |
|  | margarate (17:0) | 1.60 |
|  | 10-heptadecenoate (17:1n7) | 2.01 |
|  | stearate (18:0) | 1.31 |
|  | oleate/vaccenate (18:1) | 1.92 |
|  | nonadecanoate (19:0) | 2.35 |
|  | 10-nonadecenoate (19:1n9) | 2.18 |
|  | arachidate (20:0) | 1.44 |
|  | eicosenoate (20:1) | 1.46 |
|  | erucate (22:1n9) | 1.20 |

**Table 2.** Changes in metabolites involved in polyunsaturated fatty acid metabolism. Green: indicates significant difference (P ≤ 0.05) between the groups shown, metabolite ratio of < 1.00. Red: indicates significant difference (P ≤ 0.05) between the groups shown; metabolite ratio of ≥ 1.00.

|  |  |  |
| --- | --- | --- |
| **Polyunsaturated Fatty Acid (n3 and n6)** | heneicosapentaenoate (21:5n3) | 1.13 |
|  | hexadecadienoate (16:2n6) | 1.10 |
|  | hexadecatrienoate (16:3n3) | 0.89 |
|  | stearidonate (18:4n3) | 0.91 |
|  | eicosapentaenoate (EPA; 20:5n3) | 1.11 |
|  | docosapentaenoate (n3 DPA; 22:5n3) | 1.33 |
|  | docosahexaenoate (DHA; 22:6n3) | 1.71 |
|  | docosatrienoate (22:3n3) | 1.00 |
|  | nisinate (24:6n3) | 0.76 |
|  | linoleate (18:2n6) | 1.57 |
|  | linolenate [alpha or gamma; (18:3n3 or 6)] | 1.46 |
|  | dihomo-linolenate (20:3n3 or n6) | 1.54 |
|  | arachidonate (20:4n6) | 1.30 |
|  | docosapentaenoate (n6 DPA; 22:5n6) | 1.48 |
|  | docosadienoate (22:2n6) | 1.23 |
|  | dihomo-linoleate (20:2n6) | 1.51 |
|  | linoelaidate (tr 18:2n6) | 1.50 |
|  | mead acid (20:3n9) | 1.54 |
|  | docosatrienoate (22:3n6)* | 1.24 |

**Table 3.**
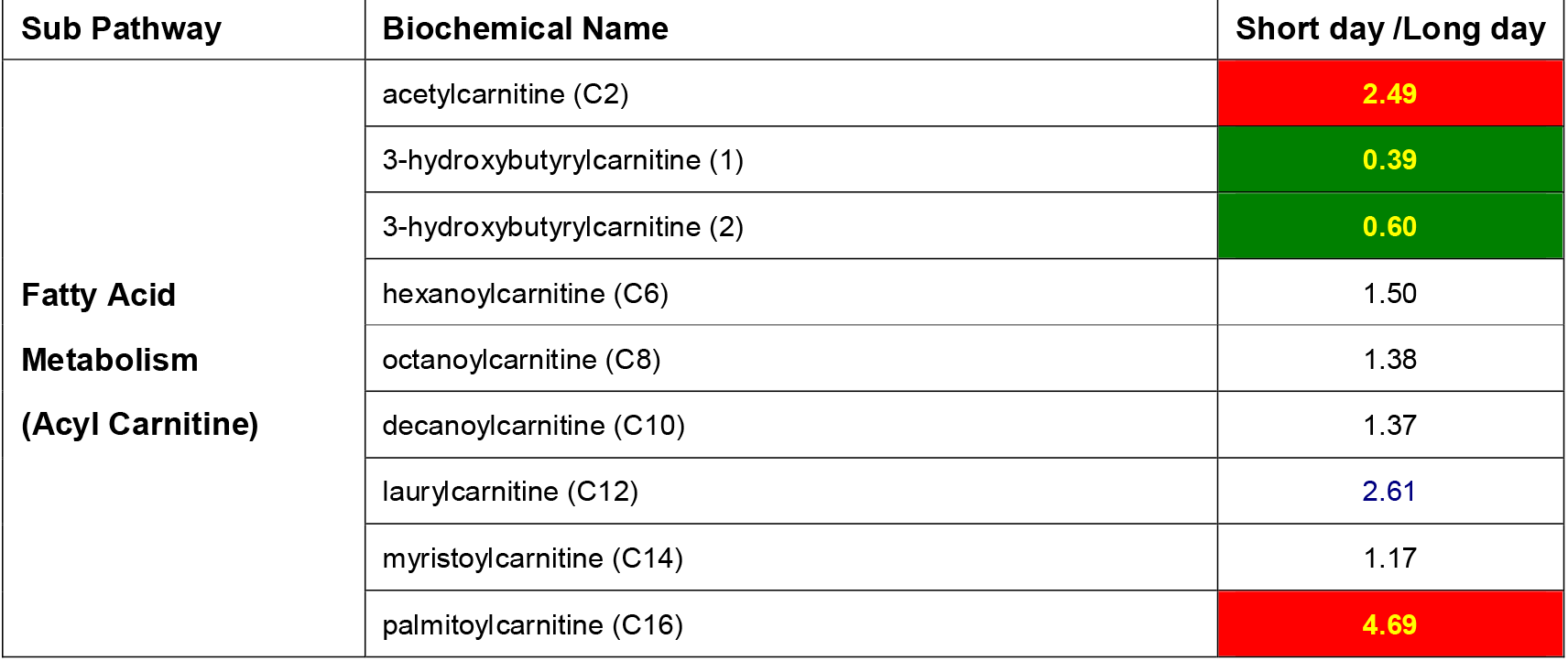
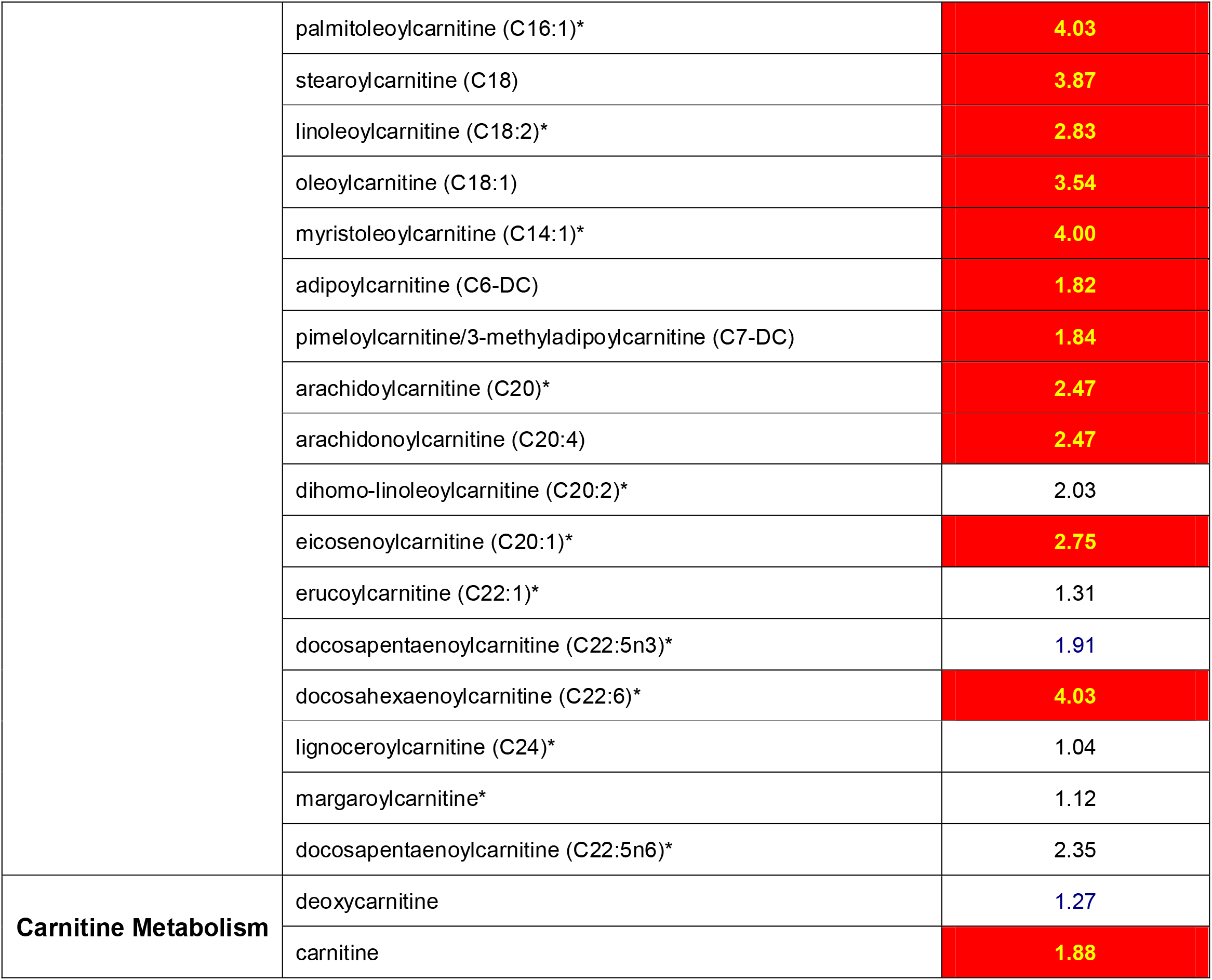
Changes in metabolites involved in carnitine metabolism. Green: indicates significant difference (P ≤ 0.05) between the groups shown, metabolite ratio of < 1.00. Red: indicates significant difference (P ≤ 0.05) between the groups shown; metabolite ratio of ≥ 1.00.

### A medaka model of fatty liver disease is efficiently established through administration of a high-fat diet under short-day conditions

Thus far, we have established medaka models of fatty liver disease through administration of a high-fat diet, but these models employed long-day conditions using photoperiods varying between L:D=14:10 and L:D=12:12. By using transparent medaka whose fatty degeneration of the liver can be observed with the naked eye, comparisons between findings under long- and short-day conditions were carried out on a high-fat diet group. Although there was no significant difference in body weights (Fig.4A), the medakas’ livers showed larger amounts of fat deposition after 14 days of breeding under short-day conditions, and marked fat deposition was observed after 28 days (Fig.4B-E). The medakas’ livers were isolated and stained with HE and fat deposition was more abundant under short-day conditions (Fig.4F). Oil-red-O staining also showed a larger amount of fat deposition under short-day conditions (Fig.4G).

**Fig. 4.**
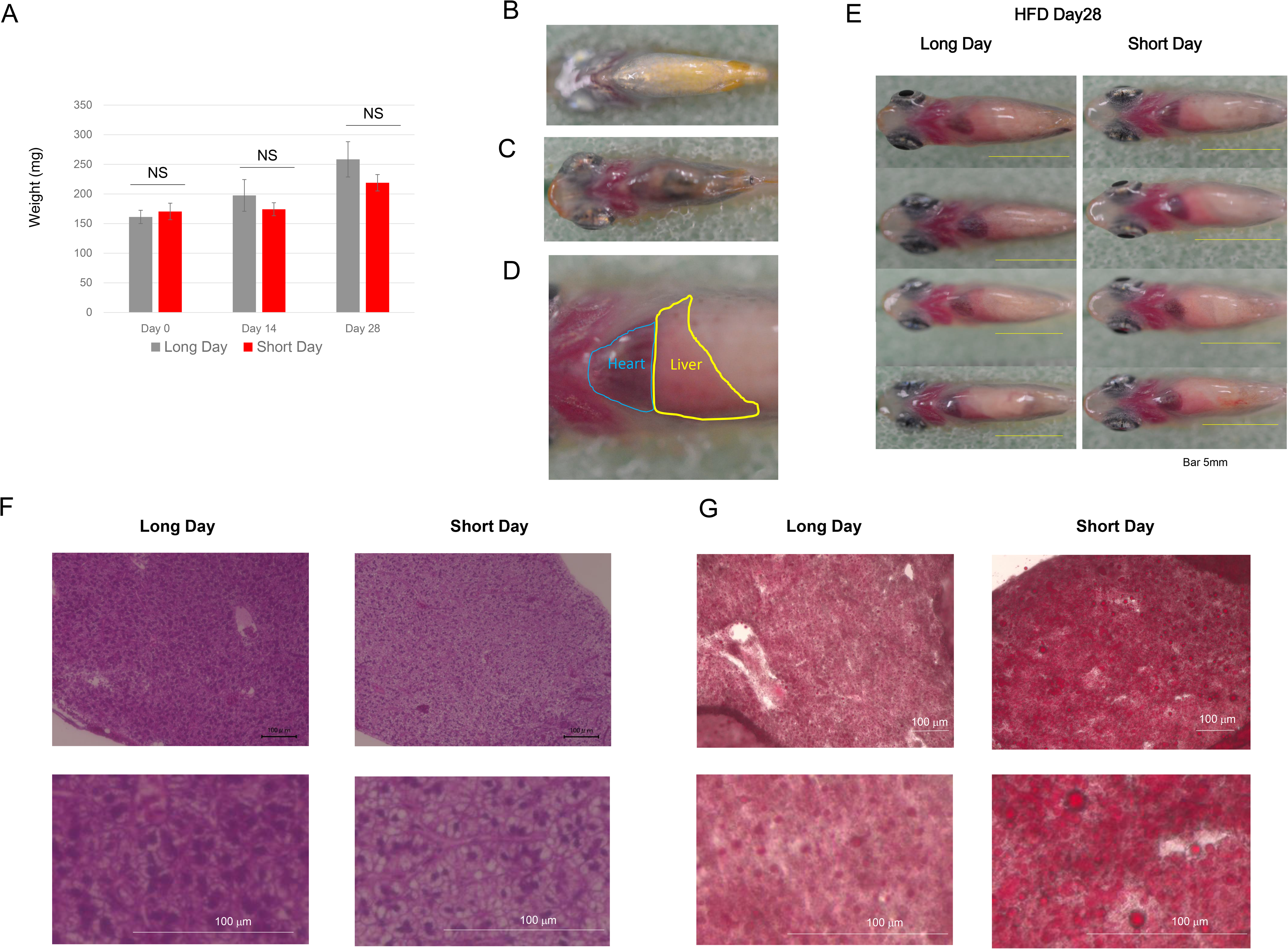
Evaluation of hepatic steatosis using a transparent medaka model fed with a high-fat diet. (A) Changes in the body weights of medaka on day 28 after exposure to variations in photoperiod. (B) WT medaka (cab strain). The liver is invisible. (C) T5 transparent medaka fed with a normal diet. The heart and the liver are visible. (D) Magnified photograph of the heart and liver of T5 transparent medaka fed with a high-fat diet. (E) T5 transparent medaka on day 28 after exposure to variations of photoperiod. Left: long day; right: short day (F) HE-stained image of the liver on day 28 after exposure to variations in photoperiod. (G) Oil-red-O-stained image of the liver on day 28 after exposure to variations of photoperiod. Data are represented as mean ± SD. Statistical significance was calculated using t-test. * indicates significant difference (P ≤ 0.05)

## Discussion

Organisms living in non-tropical regions are sensitive to seasonal variations because of their need to live through winter, during which food is scarce. In mammals, light-related information is known to be acquired from the retina and conveyed through the mediation of melatonin (Carter and Goldman, 1983). As for fish, recent reports have shown that coronet cells in the saccus vasculosus perceive changes in daylight hours and act as “seasonal sensors” to control breeding activities (Nakane et al., 2013). Particularly, amphibians and fish living in temperate climates are known to accumulate fat as energy storage during the winter. In amphibians, research studies using frogs for the study of liver metabolism and its seasonal variations have reported increases in liver weight and glycogen content from autumn to winter (Smith, 1950). Seasonal variations regarding fatty acids in little tuna (*Euthynnus alletteratus*) have been analyzed, and fatty acids in the liver reportedly increased in winter (Selmi et al., 2008). Those reports included the influence of decreased photoperiod and water temperature, but low temperature only has been reportedly associated with a larger deposition of fat in the liver of salmon (Ruyter, 2006). Previous reports regarding medaka have shown that when the temperature was constant and only the photoperiod varied, changes occurred in the expression of the long noncoding RNA that regulates adaptive behaviors to seasonal environmental changes (Nakayama et al., 2019); however, there has been no detailed analysis of the effect of photoperiod variations alone on hepatic lipid metabolism.

In our study, water temperature was kept constant and only photoperiod was varied. When fish were fed a normal diet under short-day conditions, there was no statistically significant difference in body weight, but a larger amount of fat accumulated in the liver. In addition, metabolomic analysis showed an increase in saturated long-chain fatty acids, indicating that fat accumulated in the liver more efficiently under short-day conditions. However, there was virtually no increase in PUFA under short-day conditions. Low temperatures during the winter are known to increase the degree of unsaturation of fatty acids to help maintain cell membrane fluidity (Xue et al., 2012). Long-chain PUFA in its polar lipids have been reportedly involved in the high adaptability of ayu (*Plecoglossus altivelis*) to low temperatures, and cold acclimatization has been reported to induce the activity of acyl CoA D9 desaturase in carp (*Cyprinus carpio*) liver (Trueman et al., 2000). In our study, the temperature was kept constant at 26 °C; therefore, variations in photoperiod alone did not increase the proportion of PUFA. In addition, an increase in various types of acylcarnitines suggested a decrease in mitochondrial β-oxidation, which may have been involved in the accumulation of lipids.

Further, real-time PCR showed decreased expression of the fat accumulation-associated gene PPARα under short-day conditions, which supported the hypothesis that mitochondrial β-oxidation had decreased. PPARα target genes include peroxisomal and mitochondrial fatty acid β-oxidation enzymes, proteins involved in the intracellular transport of fatty acids and the transmembrane transport of fatty acids (Motojima et al., 1998). PPARα is said to be involved in fatty acid catabolism; its expression has been found to follow a diurnal pattern and to be involved in torpor/circadian rhythm (Ishida, 2009).

As for changes related to energy metabolism, the glycolytic pathway showed no major change, but a decrease was observed in the metabolites of the TCA cycle (succinate, fumarate, malate), which suggested that citrate may have been used for fatty acid synthesis. As for interesting changes in IPA, levels of taurine increased. Taurine has been reported to have antioxidant properties (Atmaca, 2004), and serum levels of taurine in dolphins (*Tursiops truncates*) have also been reported to increase in the winter (Miyaji et al., 2012). In addition, the decrease in GSH/GSSG (glutathione disulphide) suggested enhanced oxidative stress due to accumulation of large amounts of lipids in the liver. In our study, the temperature was maintained at 26 °C and only changes in photoperiod were applied, but in the future, detailed analyses are warranted to determine the extent of the effect of temperature on hepatic fat metabolism, and whether a synergistic effect exists between photoperiod and temperature.

The lean season (namely the period during which fish have less fat in their bodies) typically coincides with egg-laying season, and consumption of the stored fat associated with gonad development is believed to affect the loss of body fat. Thus, since medakas do not carry out reproductive activities under short-day conditions, this may have been the reason why a larger amount of lipids was stored in their livers. In addition to affecting reproductive activities, various metabolic changes are believed to occur, for example, the thyroid hormones of frogs (*Rana perezi*) have been found to display seasonal changes (Gancedo et al., 1995). In the future, the mechanism that starts with the perception of photoperiod length and leads to hepatic steatosis will need to be examined, as well as its association with other organs, such as the brain, muscles and fat tissues.

Many patients currently suffer from fatty liver disease; their condition is likely to progress towards NASH, cirrhosis and liver cancer, and the risk of developing various lifestyle-related diseases is also higher. Therefore, new models aimed at conducting detailed analyses of the underlying mechanisms are needed. We have previously evaluated rats and mouse models of hepatic steatosis, and in our previous studies using medaka, which is a small-size fish species suitable for drug screening, we have also conducted detailed analysis of changes in metabolites associated with hepatic steatosis, and examined non-invasive evaluation methods. The medaka model of fatty liver disease used at that time bred under long-day conditions (14:10), and the study was carried out without paying particularly careful attention to photoperiod. In our current medaka model of fatty liver disease induced by administration of a high-fat diet, accumulation of a larger amount of fat was found under short-day conditions, demonstrating that induction of fatty liver disease could be achieved more efficiently under short-day conditions. In the future, additional studies will be needed to examine the extent of the effect of temperature, and to determine whether the latter exerts a synergistic effect with photoperiod in the development of hepatic steatosis.

Finally, regarding whether increased fat accumulation in the liver is also likely to occur in humans under short-day conditions, it has been pointed out that responses to seasonal changes have most likely decreased because modern humans use lighting at night. However, seasonal variations in human serum lipid levels have previously been assessed, and low-density lipoproteins and hepatic lipase (HL) have shown lowest levels in the summer, whereas high-density lipoproteins (cHDL) and lipoproteins (LPL) have shown lowest levels in the winter. These findings demonstrated for the first time that in physiological conditions, plasma LPL and HL activities and lipids followed seasonal rhythms (Cambras et al., 2017).

Our study revealed that even in the absence of temperature variations, more fatty acids accumulated in the liver under short-day than long-day conditions, and accumulation of acylcarnitine and unsaturated long-chain fatty acids was found under short-day conditions. Further, we were able to create steatotic medaka more efficiently when a high-fat diet was administered under short-day conditions. Conducting a study on the relationship between seasonal changes and hepatic steatosis will be of crucial importance in the future as incidences of lifestyle-diseases increase, and the findings of our study will be useful for research studies examining the relationship between photoperiod and diseases such as hepatic steatosis and NASH.

## Materials and Methods

### Experimental Model

The inbred medaka strain (Kyoto-Cab) was used in this study (Furutani-Seiki et al., 2004). Six-to ten-month-old Himedaka strain Cab (an orange-red variety of medaka) fish were used for the metabolome analysis. The fish in a given tank received a daily ration of 20 mg/fish of the diet prescribed for that group, an amount that was consumed completely within 14 h. Female 6-month-old transparent medaka (T5 strain) kindly provided by Dr. Shima (Shimada and Shima, 2001; Shimada and Shima, 2004) were used in the experiments where the progress of fatty liver was assessed. All fish were maintained in accordance with the Animal Care Guidelines of Yamaguchi University (Yamaguchi, Japan) (approval number is 21-038). During experiments, fish were kept in plastic tanks covered with plastic covers and illuminated with fluorescent light from 8:00–20:00. The tank water temperature was maintained at 26 °C.

### Diets

The proportions of protein, fat and carbohydrate, as well as the fatty acid compositions, of the control and high-fat diets used in this study were previously reported (Matsumoto et al., 2010). The energy content of the control diet (Hikari Crest; Kyorin Co. Ltd, Hyogo, Japan) was 3.3 kcal/g, with 25.3% of calories from fat, 62.5% of calories from protein, and 13.8% of calories from carbohydrate. The energy content of the high-fat diet (HFD32; CLEA Japan Inc., Tokyo, Japan) was 5.1 kcal/g, with 56.7% of calories from fat, 20.1% of calories from protein, and 23.2% of calories from carbohydrate.

### Twenty-four hour behavior monitoring

Using the Chronobiology Kit (Stanford Software Systems, Naalehu, HI, USA), ten medaka were placed in one water tank, and the amount of physical exercise and periodicity of their activity were observed per 24 h. The experiment was conducted over a 10-day period, and the results were displayed as actograms. Light-dark conditions were as follows: breeding was carried out using L:D=14:10 (h) (light phase: 8:00□22:00, dark phase: 22:00□8:00) for 10 days, after which the conditions were changed to L:D=10:14 (h) (light phase: 10:00□20:00, dark phase: 20:00□10:00) and evaluations were conducted for a 20-day period under 10/14 short-day conditions (L:D=10:14).

### Adjustment of the number of hours of sunlight

Ten medaka were placed in one water tank and bred using L:D=14:10 (h) under long-day conditions and L:D=10:14 (h) under short-day conditions. Adult medaka which had been bred under L:D = 14:10 for 28 days or longer were reared separately in two groups, namely the short-day group and the long-day group (Fig.1B). The medaka were fed with a normal diet, and their weights measured and livers isolated 28 days later.

### Real-time-PCR method

Using the StepOnePlus Real-time PCR System (Applied Biosystems, Waltham, MA, USA) and SYBR Green, variations in the expression levels of 18S rRNA and PPARα were analyzed. For RT-PCR analysis, primers with relatively good amplification efficiencies and without non-specific amplification in the dissociation curves were selected. The base sequences of the primers used in the reactions were as follows.

PPARα Forward 5′-AGGGTTGCAAGGGTTTCTTT-3′
PPARα Reverse 5′-AGCTTCAGCTTCTCCGACTG-3′
EF1α Forward 5′-AACACTCCTTGAAGCTCTTG-3′
EF1α Reverse 5′-GACAGGGACAGTTCCAATAC-3′

### Metabolomic analysis

Metabolomic and statistical analyses were conducted at Metabolon, Inc. as described previously (Shin et al., 2014). Briefly, cell pellets were subjected to methanol extraction and then split into aliquots for analysis by ultrahigh performance liquid chromatography/mass spectrometry (UHPLC/MS) in the positive, negative or polar ion mode and by gas chromatography/mass spectrometry (GC/MS). Metabolites were identified by automated comparison of ion features to a reference library of chemical standards followed by visual inspection for quality control. To identify the affected metabolic pathways, a proof-of-knowledge based Ingenuity Pathway Systems (IPA, Redwood City, CA, USA) analysis was performed.

### Statistics

To determine the statistical significance of the real-time PCR analysis, t-tests were used; P < 0.05 was considered significant. To determine the statistical significance of the metabolome analysis, Welsh’s two-factor t-tests were performed in Array Studio (Omicsoft Corporation, Cary, NC, USA) or “R” to compare protein-normalized data between experimental groups; P < 0.05 was considered significant.

## Acknowledgments

We would like to thank Ms. Mariko Yamada, Ms. Risa Mochizuki and Ms. Kumie Ota for their technical assistance.

